# Computational simulations reveal the binding dynamics between human ACE2 and the receptor binding domain of SARS-CoV-2 spike protein

**DOI:** 10.1101/2020.03.24.005561

**Authors:** Cecylia S. Lupala, Xuanxuan Li, Jian Lei, Hong Chen, Jianxun Qi, Haiguang Liu, Xiao-dong Su

**Author notes:** Corresponding authors: Haiguang Liu, and Xiao-dong Su.

## Abstract

A novel coronavirus (the SARS-CoV-2) has been identified in January 2020 as the causal pathogen for COVID-19 pneumonia, an outbreak started near the end of 2019 in Wuhan, China. The SARS-CoV-2 was found to be closely related to the SARS-CoV, based on the genomic analysis. The Angiotensin converting enzyme 2 protein (ACE2) utilized by the SARS-CoV as a receptor was found to facilitate the infection of SARS-CoV-2, initiated by the binding of the spike protein to the human ACE2. Using homology modeling and molecular dynamics (MD) simulation methods, we report here the detailed structure of the ACE2 in complex with the receptor binding domain (RBD) of the SARS-CoV-2 spike protein. The predicted model is highly consistent with the experimentally determined complex structures. Besides the binding interface reported in the crystal structures, new plausible binding poses were revealed from all-atom MD simulations. The simulation data helped identify critical residues at the complex interface and provided more details about the interactions between the SARS-CoV-2 RBD and human ACE2. Two mutants mimicking rat ACE2 were modeled to study the mutation effects on RBD binding to ACE2. The simulations showed that the N-terminal helix and the K353 are very important for the tight binding of the complex, the mutants were found to alter the binding modes of the CoV2-RBD to ACE2.

## Introduction

The outbreak of a new type of severe pneumonia COVID-19 started in December 2019^1^ has been going on world-wide, and caused over 15,000 fatalities, infected more than 350,000 individuals globally. Although the earlier infected cases were mainly found in China before March, 2020, particularly in Hubei Province, the confirmed COVID-19 cases have been reported in more than 160 countries or territories by the end of March, 2020, and still increasing rapidly. One urgent desire in coping with this global crisis is to develop or discover drugs that can treat the diseases caused by the novel coronavirus, the SARS-CoV-2 (also known as 2019-nCoV)^2^. According to the genome comparative studies, the SARS-CoV-2 belongs to the genus beta-coronavirus, with nucleotide sequence identity of about 96% compared to the closest bat coronavirus RaTG13, about 89% compared to two other bat SARS-like viruses (Bat-SL-CoVZC45 & Bat-SL-CoVZXC21), and 79% compared to the SARS-CoV^3,4^ Furthermore, the SARS-CoV-2 spike protein has a protein sequence identity of 73% for the receptor binding domain (RBD) with the SARS-CoV RBD (denoted as SARS-RBD in the following). The SARS-CoV and SARS-CoV-2 both utilize the human Angiotensin converting enzyme 2 protein (ACE2) to initiate the spike protein binding and facilitate the fusion to host cells^5–9^. The 193-residue RBD of the SARS-CoV spike protein has been found to be sufficient to bind the human ACE2^6^. Based on this fact, the RBD of SARS-CoV-2 becomes a critical protein target for drug development to treat the COVID-19. When this study was started, neither the crystal structure of the SARS-CoV-2 spike protein nor the RBD segment were determined, so the homology modeling approach was applied to construct the model of the SARS-CoV-2 spike RBD in complex with the human ACE2 binding domain (denoted as CoV2-RBD/ACE2 in the following). Similar approach has been applied to predict the complex structure and estimate the binding energies^3^. Because of the high sequence similarity between CoV2-RBD and SARS-RBD, the predicted structure was found to be highly consistent with the resolved crystal structures^10^ (see another crystal structure at http://nmdc.cn/nCoVentry:NMDCS0000001, PDBID: 6LZG). These structures laid the foundation for the dynamics investigation of the CoV2-RBD/ACE2 complex using computational simulation method. The predicted CoV2-RBD/ACE2 model was subjected to all-atom molecular dynamics (MD) simulations to study the binding interactions.

Although the crystal structure and the predicted model of the CoV2-RBD/ACE2 complex provide important information about the binding interactions at the molecular interfaces, MD simulations can extend the knowledge to a dynamics regime in a fully solvated environment. The importance of the ACE2 residues was investigated by simulating the complexes with ACE2 mutants, in which partial dissociation from the ACE2 was observed within 500 ns simulations. The control simulations of the SARS-RBD/ACE2 complexes allowed the detailed comparison in receptor binding for the two different types of viruses. The results showed that the wild type CoV2-RBD/ACE2 complex is stable in 500 ns simulations, especially in the well-defined binding interface. On the other hand, the mutations on the helix-1 or K353 of the ACE2 can alter the binding, revealing new binding poses with reduced contacts compared to those in the crystal structures. The analysis of the interaction energy showed that the binding is enhanced by adjusting conformations to form more favorable interactions as the simulation progressed, consistent with the increased hydrogen bonding patterns. Furthermore, the analysis also showed that SARS-RBD and CoV2-RBD have comparable binding affinities to the ACE2, with the former slightly stronger than the latter in terms of molecular mechanics energies. The dynamic information obtained by this study shall be useful in understanding SARS-CoV-2 host interaction and for designing inhibitors to block CoV2-RBD binding.

## Methods

### The computer model of the SARS-CoV-2 spike RBD in complex with human ACE2

The spike RBD of SARS-CoV-2 (GenBank: MN908947^2^) comprises Cys336-Gly526 residues according to the sequence homology analysis with SARS-CoV spike RBD (see Figure S1). The predicted three-dimensional structure model of these residues was obtained with the SWISS-MODEL server^11^. This predicted SARS-CoV-2 RBD model was subsequently superimposed into the X-ray structure of SARS-CoV RBD in complex with human ACE2 (PDB code:2AJF, Chain D^7^). Finally, the computer model of SARS-CoV-2 RBD with human ACE2 (CoV2-RBD/ACE2) was obtained for further simulations and analysis.

### Mutant preparation

Based on the analysis of the predicted model, sequence alignment, and literature survey, two other systems containing mutations in the human ACE2 were prepared and subject to MD simulation studies. The mutant construct is based on the fact that Rat ACE2 markedly diminishes interactions with SARS spike protein^12^, and it was proposed that the rat ACE2 likely has reduced binding affinity to the CoV2-RBD^13^. To investigate the roles of critical residues on the ACE2, we created two mutants of the human ACE2 (see Table 1): (1) mutant *mut_h1*, with the ACE2 N-terminal (residue 19-40) mutated to the residues of rat ACE2; and (2) mutant *K353H,* in which the highly conserved K353 was mutated to histidine (the corresponding amino acid in rat and mouse ACE2 proteins). To focus on the impact of these two binding sites, the rest of the ACE2 were kept to be the same as human ACE2.

**Table 1.** The ACE2 mutants to mimic the Rat ACE2.

| Mutant name | Mutations on ACE2 | Remark |
| --- | --- | --- |
| mut_h1 | T20L, Q24K, K26E, T27S, D30N, H34Q, F40S | L20, N30, Q34, and S40 are conserved between rat and mouse. |
| K353H | K353H | H353 is conserved between rat and mouse |

### Molecular Dynamics Simulation and Analysis

The predicted model of CoV2-RBD/ACE2 complex was used as the starting models for MD simulations. The spike protein RBD domain is composed of 180 residues (323-502), while the ACE2 protein contains 597 residues (19-615) from the N-terminal domain. The simulation parameterization and equilibration were prepared for complex structures including the mutant systems, using the CHARMM-GUI webserver^14^ Each system was solvated in TIP3P water and sodium chloride ions were added to neutralize the systems to a salt concentration of 150 mM. Approximately, each system was composed of about 220,000 atoms that were parametrized with the CHARMM36 force field^15^.

After energy minimization using the steepest descent algorithm, each system was equilibrated at human body temperature 310.15 K, which was maintained by Nose-Hoover scheme with 1.0 ps coupling constant in the NVT ensemble (constant volume and temperature) for 125.0 ps under periodic boundary conditions with harmonic restraint forces applied to the complex molecules (400 kJ mol^-1^ nm^-2^ on backbone and 40 kJ mol^-1^ nm^-1^ on the side chains)^16,17^. In the subsequent step, the harmonic restraints were removed and the NPT ensembles were simulated at one atmosphere pressure (10^5^ Pa) and body temperature. The pressure was maintained by isotropic Parrinello-Rahman barostat^18^, with a compressibility of 4.5 □ × □ 10^-5^ bar^-1^ and coupling time constant of 5.0 □ ps. The simulation trajectories were propagated to 500 nanoseconds using the GROMACS 5.1.2 package^19^

Three independent trajectories starting from random velocities based on Maxwell distributions were simulated for both CoV2-RBD/ACE2 and SARS-RBD/ACE2 complex systems in their wild types. In all simulations, a time step of 2.0 fs was used and the PME (particle mesh Ewald)^20^ was applied for long-range electrostatic interactions beyond 12.0 Å. The van der Waals interactions were evaluated within the distance cutoff of 12.0 Å. Hydrogen atoms were constrained using the LINCS algorithm^21^.

The human ACE2 mutants in complex with CoV2-RBD were constructed as described previously. Each mutant complex model was simulated in two independent trajectories. Furthermore, as the crystal structure of the CoV2-RBD/ACE2 complex became available, two additional simulations were carried out using the crystal structure as the starting model to cross-validate the simulation results based on the homology model. Each trajectory was propagated to 500 ns by following the same protocol as the wild type CoV2-RBD/ACE2 complex simulations.

Analyses were carried out with tools in GROMACS (rmsd, rmsf, energy, and pairdist) to examine the system properties, including the overall stability, local residue and general structure fluctuations through the simulations. The *g_mmpbsa* program^22^ was applied to extract the molecular mechanics energy E_MM_ (Lennard Jones and electrostatic interactions) between ACE2 and the RBD of spike proteins. VMD and Chimera were applied to analyze the hydrogen bonds, molecular binding interface, water distributions, visualization, and rending model images^23,24^

## Results

The homology structure of the CoV2-RBD/ACE2 was compared to both the SARS-RBD/ACE2 crystal structure and the newly resolved crystal structure of the CoV2-RBD/ACE2. The results indicated that the homology model is accurate, especially at the binding interface. The MD simulations further refined the side chain orientations to improve the model quality. The simulation data confirmed the highly stable binding between the CoV2-RBD and the ACE2, in spite of the conformational changes of the ACE2. The relative movement between the CoV2-RBD and the ACE2 mainly exhibited as a swinging motion pivoted at the binding interface. Simulations also revealed the roles of water molecules in the binding of the RBD to the ACE2 receptor. The MD simulation of complex with ACE2 mutants suggested that mutation to the ACE2 helix-1 and the K353 can alter the binding modes and binding affinity.

### 1. Homology modeling and comparison to the SARS-RBD/ACE2 complex

The predicted CoV2-RBD/ACE2 complex structure is highly similar to the SARS-RBD/ACE2, as shown in Figure 1. The RBD domain has an RMSD of 0.99 Å for the aligned residues (1.53 Å for all 174 residue pairs), indicating that the homology model of the CoV2-RBD is in good agreement with the SARS-RBD. For ACE2 residues near the binding interface (within 4.0 Å of the RBD), the RMSD is smaller than 0.43 Å compared to the SARS-RBD/ACE2 complex. The superposed structures revealed that the RBD/ACE2 interfaces are almost identical in two complexes(Figure 1c).

**Figure 1.**
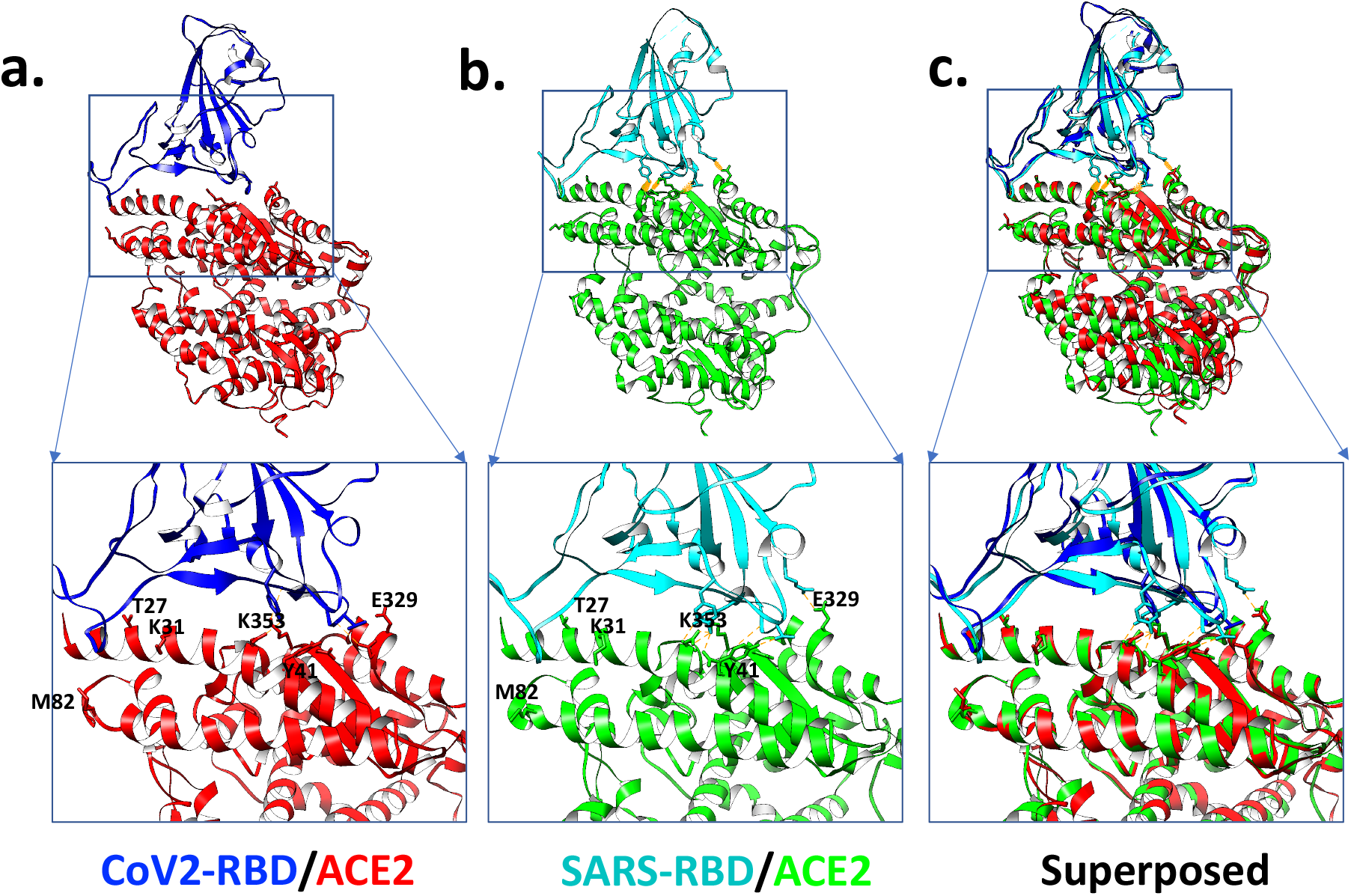
The predicted structure and the binding interface. (**a**) Predicted model of the CoV2-RBD/ACE2; (**b**) Crystal structure of the SARS-RBD/ACE2 (PDB: 2AJF); (**c**) Superposed models. The binding interface is indicated in the boxed region.

In a retrospective comparison, the homology model was superposed to a newly resolved crystal structure (PDB ID: 6LZG; see Figure 2d for a detailed comparison at the interface). The results indicate that the homology model is very accurate, especially for the binding interface. The residues near the CoV2-RBD/ACE2 interface (defined as the combined set of ACE2 residues within 4.0 Å of RBD and the RBD residues within 4.0 Å of the ACE2) exhibited a difference of 0.43 Å RMSD, which is comparable to the difference between the two independently reported crystal models (an RMSD of 0.25 Å for the same comparison). The RMSD is about 0.77 Å for residues in an extended region within 10.0 Å of the binding interface. The RBD domain of the spike protein showed an overall RMSD values less than 1.5 Å, and the ACE2 domain with an RMSD about 2.0 Å between the predicted model and the crystal structures.

**Figure 2.**
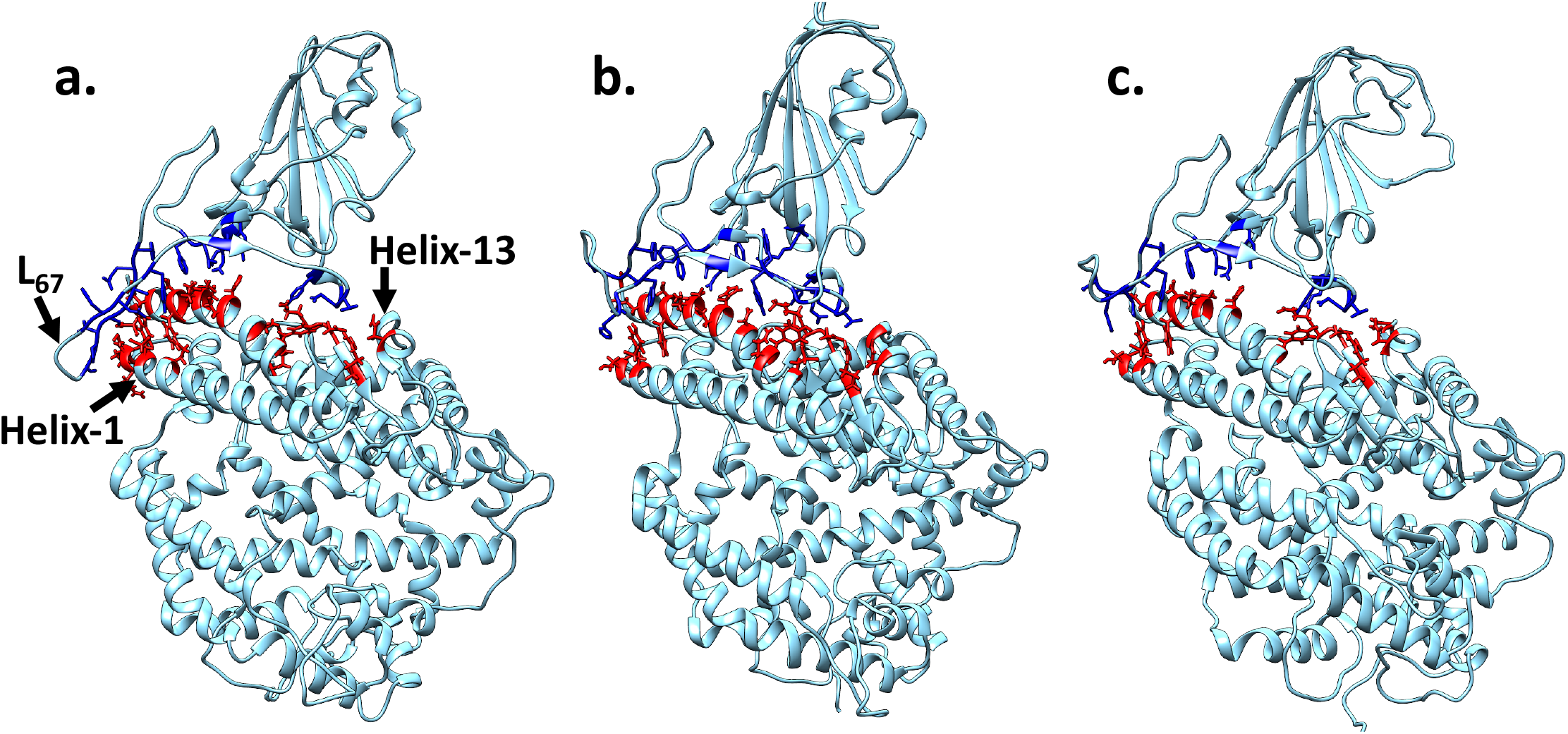
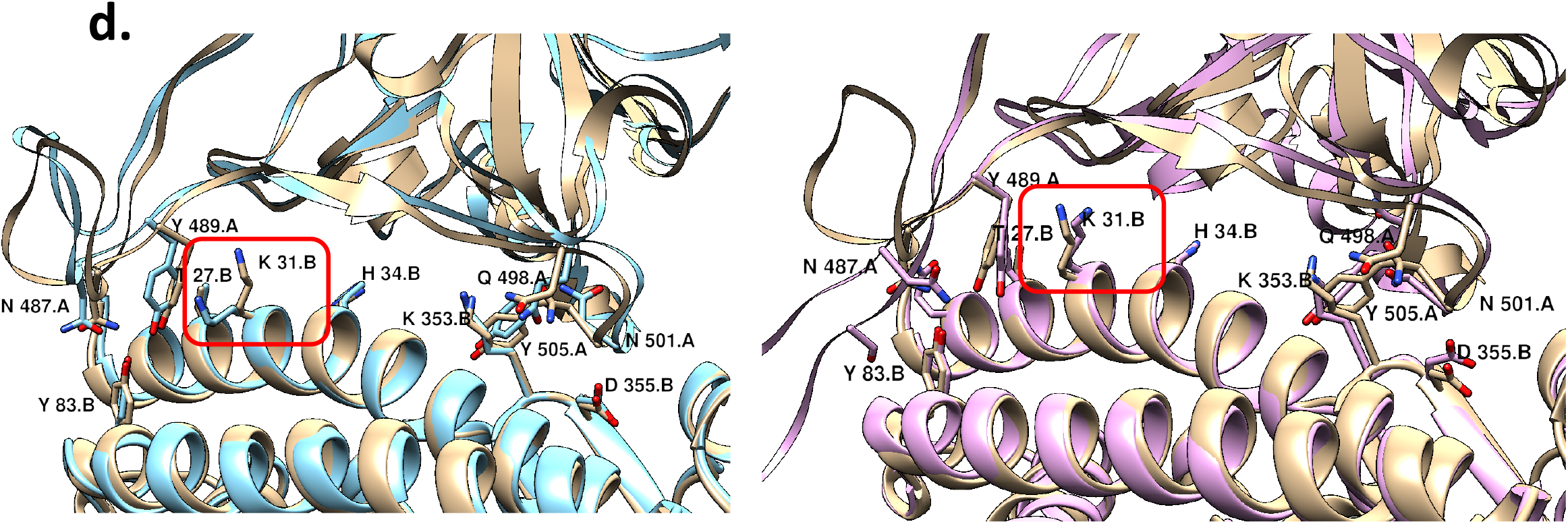
Representative structures extracted from simulation trajectories using clustering analysis. (**a**) The most populated structure from simulation trajectory#1, with the loop (L_67_, S477-G485) between β6 and β7 moved towards the ACE2 helix-1, forming additional contacts. (**b, c**) The most populated structures in the simulations #2 and #3. The complex structures and the binding interfaces are similar to the predicted model and the crystal structure. (**d**) The comparison of initial (cyan) and representative (pink) structures to the crystal structure (gold) of CoV2-RBD/ACE2. The side chain of K31 (encircled in the red boxes) was refined to the correct position during the MD simulations (right panel).

### 2. The conformational changes of the ACE2 and the CoV2-RBD

The structural comparison with respect to the starting structure revealed that MD simulations mainly sampled the structures in the conformational space near the starting model. The RMSD with respect to the starting model indicated that the sampled structures were mostly 2.5 Å to 5.0 Å compared to the starting model (Figure 3a). The largest contribution to the conformational changes in CoV2-RBD is from the loop regions, indicated by the peaks shown in the fluctuation plots (Figure 3b). The residues with well-defined secondary structures (mainly β-strands) exhibited very small conformational changes (the RMSD is about 1.0 Å including side chain atoms), while the loop regions have RMSD values between 3.7 Å to 5.4 Å. This is consistent with the residue fluctuations measured with the RMSF (root-mean-square-fluctuation) for the RBD subunit (Figure 3b). For the ACE2 receptor domain, the major conformational change is the opening/closing of the enzymatic active site, which is remote from the RBD binding interface (see Video 1). The CoV2-RBD remained bound to human ACE2, even though substantial conformational changes occurred to the ACE2 and the loop region of the RBD. This observation implies that the binding of CoV2-RBD to the ACE2 is very stable, similar with the case of the SARS-RBD that binds to both open and closed forms of human ACE2^7,25^.

**Figure 3.**
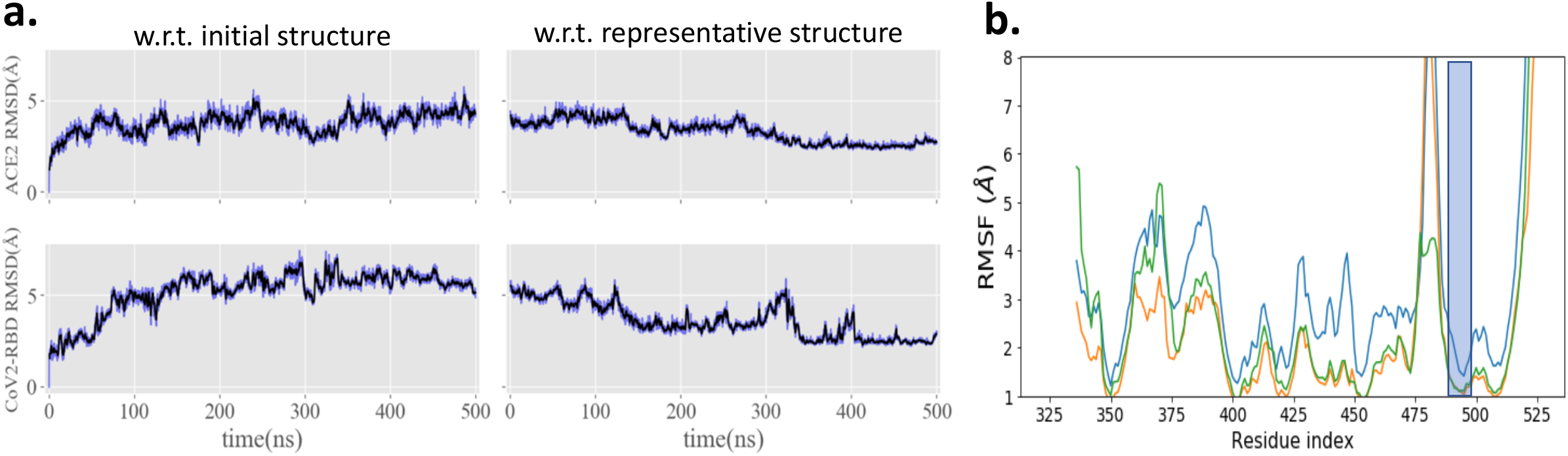
The dynamics analysis of the simulation trajectory. (**a**) The root-meansquare-deviation (RMSD) of the simulated structures compared to the initial and representative models. (**b**) The residue level fluctuations of CoV2-RBD in three simulations. The shaded area corresponds to the CoV2-RBD binding region to the ACE2.

Based on the structural similarity, a clustering analysis was carried out to identify the most populated structures, from which the representative model for each trajectory was selected (Figure 3). The CoV2-RBD/ACE2 interfaces were highlighted using blue/red colors with stick representations. The RMSD of structures in the simulations were calculated with respect to the these representative models, in order to assess the model convergence and coverage of the simulated conformations (Figure 3a, right panels). The RMSD values indicated that a large portions of the structures sampled by the MD simulations are similar to the representative model (Figure 3a, for trajectory 1, see Figure S2 for the other trajectories). Furthermore, as the simulation progressed, the conformations converged to the representative model obtained from this trajectory. This indicates that the selected structure is plausible and stable. The representative structures for the other two trajectories showed smaller deviations compared to the predicted model (Figure 2b and 2c).

In three simulations of the CoV2-RBD/ACE2 systems, the binding interface was highly stable, exhibiting very small conformational changes, especially for the interfacing residues of the ACE2 protein. The RMSD for the residues at the RBD binding interface is 0.85Å (+/-0.13Å) on average. Side chain atom positions were refined to form more favorable interactions (Figure 2d). One outstanding example is the K31 side-chain, which pointed in the wrong orientation in the predicted structure, was quickly refined to the correct orientation, consistent with the crystal structure (right panel of Figure 2d).

In terms of collective conformational changes, the CoV2-RBD/ACE2 complex showed two interesting movements: (1) the loop (L_67_) between β_6_ and β_7_ (residues between S477 and G485 in particular) of CoV2-RBD was found to expand its interactions with the N-terminal helix (the helix-1) of the ACE2 (Figure 2a), while it pointed away from the helix-1 in the predicted and the crystal structures (Figure 2d, left); (2) a tilting movement of the RBD relative to the ACE2 was observed, which can be depicted as a swinging motion with the binding interface as the pivot (see Figure 4 for an illustration). In both predicted and the crystal structures of the CoV2-RBD/ACE2 complex, the L_67_ does not form close contacts with the ACE2. The analysis of the crystal packing revealed that this loop participated in the interaction with another asymmetric unit (see Figure S3). Interestingly, the simulation data suggested that the L_67_ could move towards the ACE2 and form contacts with the helix-1. This can potentially enhance the binding, as reflected in the change of interaction energies. In the crystal structure, the C480 and C488 of the RBD are crosslinked via a disulfide bond that reduces the flexibility of the L_67_ region, limiting its access to the ACE2. On the other hand, it has been reported that the binding of SARS-RBD to ACE2 is insensitive to the redox states of the cysteines to a high extend^26^. Based on the simulation results, we hypothesize that the reduced form of C480 and C488 can also exist during the virus invasion to host cells, and the reduced cysteines can potentially enhance the binding to ACE2. In the other two simulation trajectories, we found that the L_67_ remained in conformations similar to that in the crystal structure and the cysteines (C480 and C488) were close enough for disulfide bond formation.

**Figure 4.**
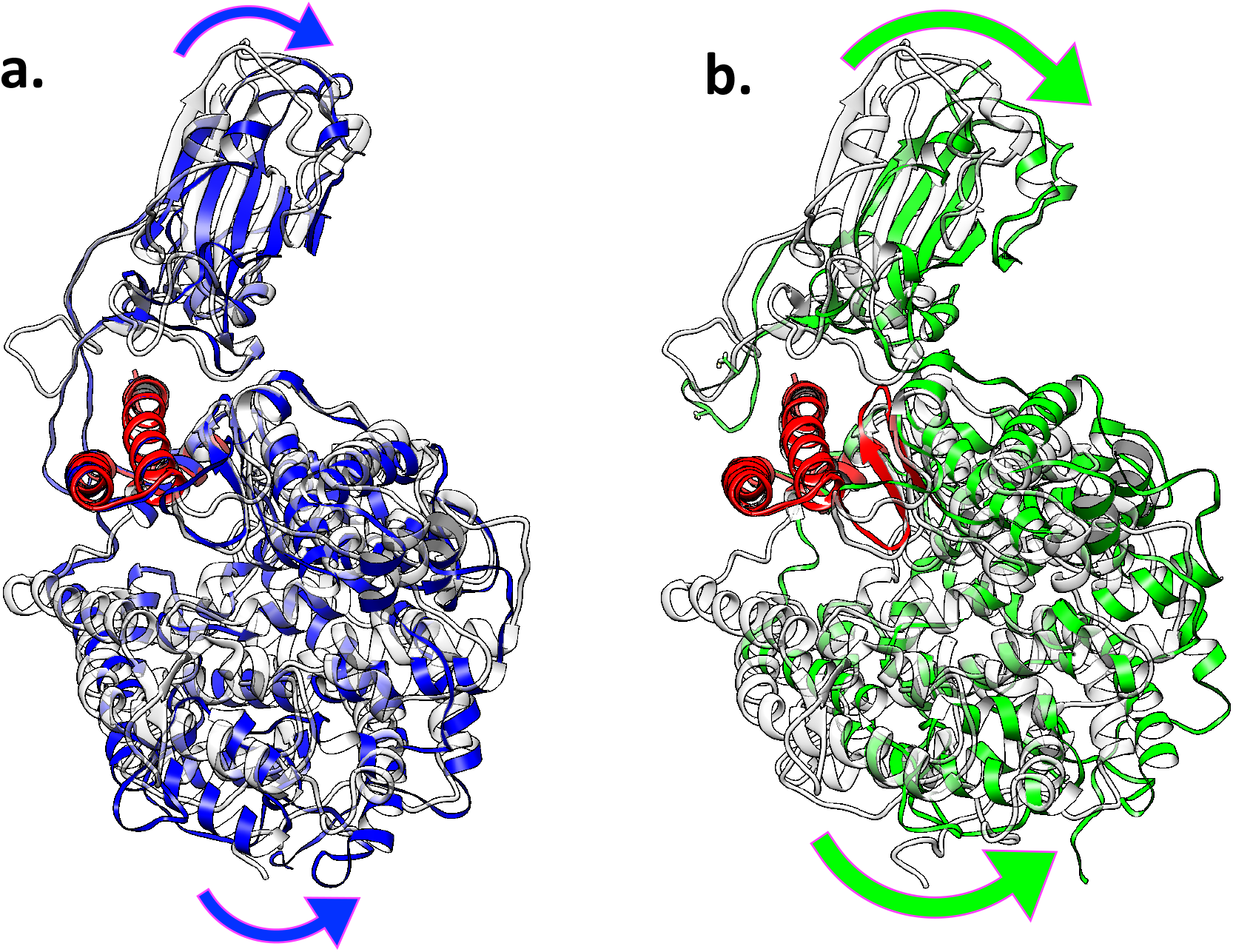
The collective motion of the CoV2-RBD/ACE2 complex. **(a)** The representative model (blue) from wild type CoV2-RBD/ACE2 simulations superposed to the initial model (silver). The arrows indicate the directions of the collective motion, and the hinge region is colored in red. **(b)** The same representation for the CoV2-RBD complexed with the ACE2 h1-mut mutant (green), and comparison between the initial model (silver) and the representative model (green) indicates a large tilting movement compared to the wild type system.

### 3. Simulations revealed detailed binding interface interactions between the CoV2-RBD and the ACE2

By examining the binding interface of CoV2-RBD and the ACE2, we found the polar and charged residues account for a large fraction, therefore the electrostatic interactions play critical roles for the complex formation. Based on the distances between the two proteins, the key residues at the binding interface were identified and summarized in Table 2 for the three representative models (see Figure 2). Majority of these residues are conserved for these three models, except that the model#1 (Figure 2a) has additional contacts to the ACE2 from residues in the L_67_ region (highlighted with green color in Table 2). As shown in Figure 2, the L_67_ remained in the starting position for the other two representative models (Figure 2b,c). The same simulations were carried out for the SARS-RBD/ACE2 complex, serving as a comparative system. Interestingly, the SARS-RBD counterpart of the L_67_ in CoV2-RBD did not form close contacts with the ACE2 in three simulations. It is worthwhile to mention that the sequence identity between CoV2-RBD and SARS-RBD is low in this loop region, suggesting the loop region might be partially responsible for the difference in the receptor binding.

**Table 2.** Contact residues between the CoV2-RBD and the ACE2. Green color denotes new interaction not observed in crystal structure.

| Model#1 |  | Model#2 |  | Model#3 |  |
| --- | --- | --- | --- | --- | --- |
| ACE2 | CoV | ACE2 | CoV | ACE2 | CoV |
| Q24 | K417 | S19 | K417 | Q24 | K417 |
| T27 | Y453 | Q24 | G446 | T27 | G446 |
| F28 | L455 | T27 | Y449 | F28 | Y449 |
| D30 | F456 | F28 | Y453 | D30 | Y453 |
| K31 | Q474 | D30 | L455 | K31 | L455 |
| H34 | A475 | K31 | F456 | H34 | F456 |
| E35 | G476 | H34 | A475 | E35 | A475 |
| E37 | S477 | E35 | F486 | E37 | F486 |
| D38 | T478 | D38 | N487 | D38 | N487 |
| Y41 | G485 | Y41 | Y489 | Y41 | Y489 |
| Q42 | F486 | Q42 | Q493 | Q42 | Q493 |
| M82 | N487 | L45 | Y495 | M82 | Y495 |
| Y83 | Y489 | M82 | G496 | Y83 | G496 |
| N330 | Q493 | Y83 | Q498 | N330 | Q498 |
| K353 | Y495 | N330 | T500 | K353 | T500 |
| G354 | G496 | K353 | N501 | G354 | N501 |
| D355 | Q498 | G354 | G502 | D355 | G502 |
| R357 | T500 | D355 | Y505 | R357 | Y505 |
|  | N501 | R357 |  |  |  |
|  | G502 |  |  |  |  |
|  | Y505 |  |  |  |  |

The hydrogen bonds between the CoV2-RBD and ACE2 were extracted using VMD program with default criteria (D-A distance cutoff=3.0 Å and D-H-A angle cutoff=20 degrees, where D,A,H are Donor atom, Acceptor atom, and the Hydrogen atom linked to the Donor atom). The numbers of hydrogen bonds are summarized in Figure 5 for the simulation trajectory#1. Because of the stringent criteria for hydrogen bonds, the numbers are smaller compared to the reported values in other studies. For a larger distance cutoff of 3.9Å, there are about 8.3 hydrogen bonds on average (Figure 5b). We observed that the number of hydrogen bonds fluctuated over time, and exhibiting an overall increasing trend indicated by the fitted lines (Figure 5). Similar trends were observed in the other simulations, suggesting that the binding became stronger as the simulation progressed (see Figure S4). Based on the statistics of three simulation trajectories, the CoV2-RBD/ACE2 complex has 2.7 hydrogen bonds between the subunits on average with stringent criteria. In comparison, the SARS-RBD/ACE2 has 3.2 hydrogen bonds on average (see Figure S4 for details). The statistics of hydrogen bonds suggest a slightly weaker binding between the CoV2-RBD and the ACE2, relative to the SARS-RBD/ACE2 complex.

**Figure 5.**
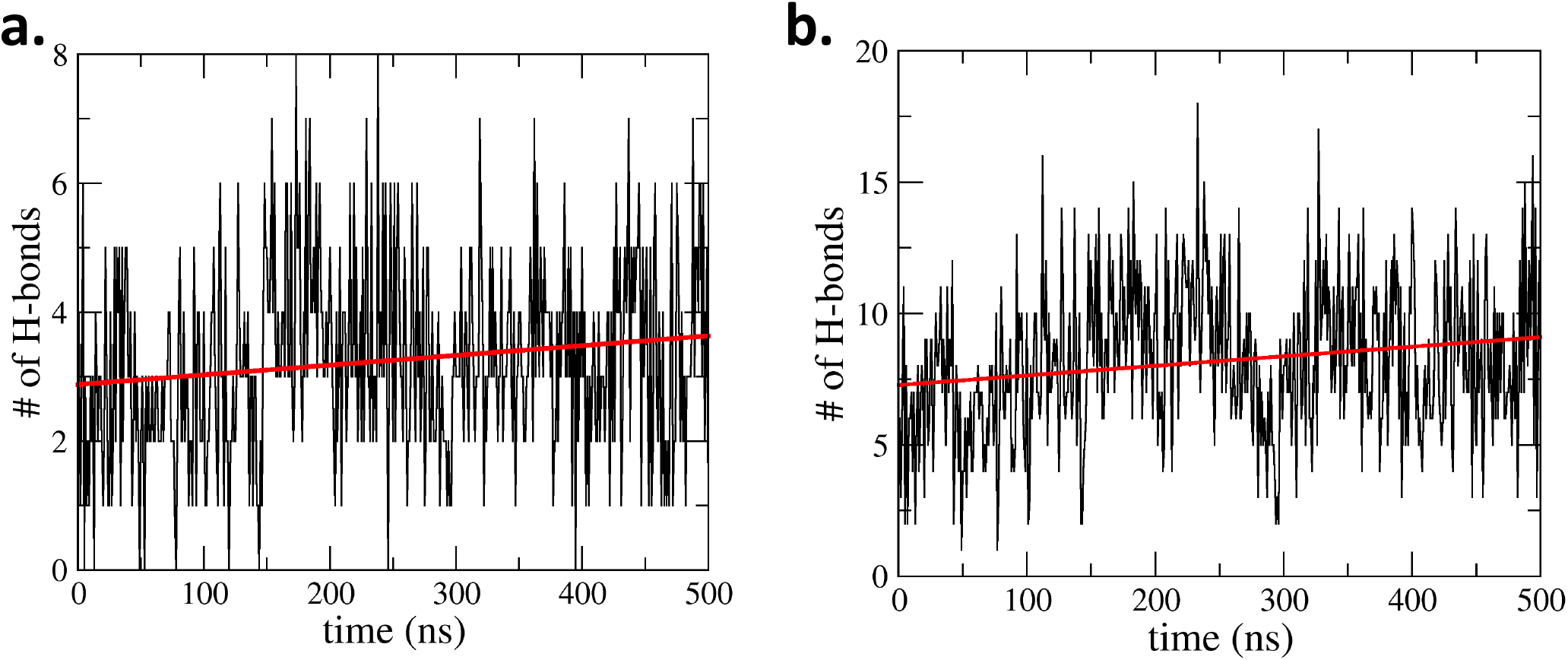
The hydrogen bonds between the CoV2-RBD and the ACE2. (**a**) The number of hydrogen bonds extracted from the simulation data using Donor-Receptor distance cutoff of 3.0 Å. (**b**) Same as (a) except that the cutoff distance was relaxed to 3.9 Å. The trend lines were obtained by least square fitting.

It is also noteworthy to point out the important roles of water molecules at the complex interface for CoV2-RBD/ACE2 complex. At any instant time, there are approximately 15 water molecules at the binding interface, simultaneously located within 2.5 Å of both the CoV2-RBD and the ACE2 (Figure 6). These water molecules can function as bridges by forming hydrogen bonds with the residues from the RBD or the ACE2. The dwelling time of water molecules at the interface can be a few nanoseconds, as revealed by the MD simulations. This results is also consistent with the crystal structure, which has 12 water molecules at the interface (Figure 6c). These discoveries emphasize the role of the water molecules, which desires detailed quantification to understand the interactions between the RBD and the ACE2.

**Figure 6.**
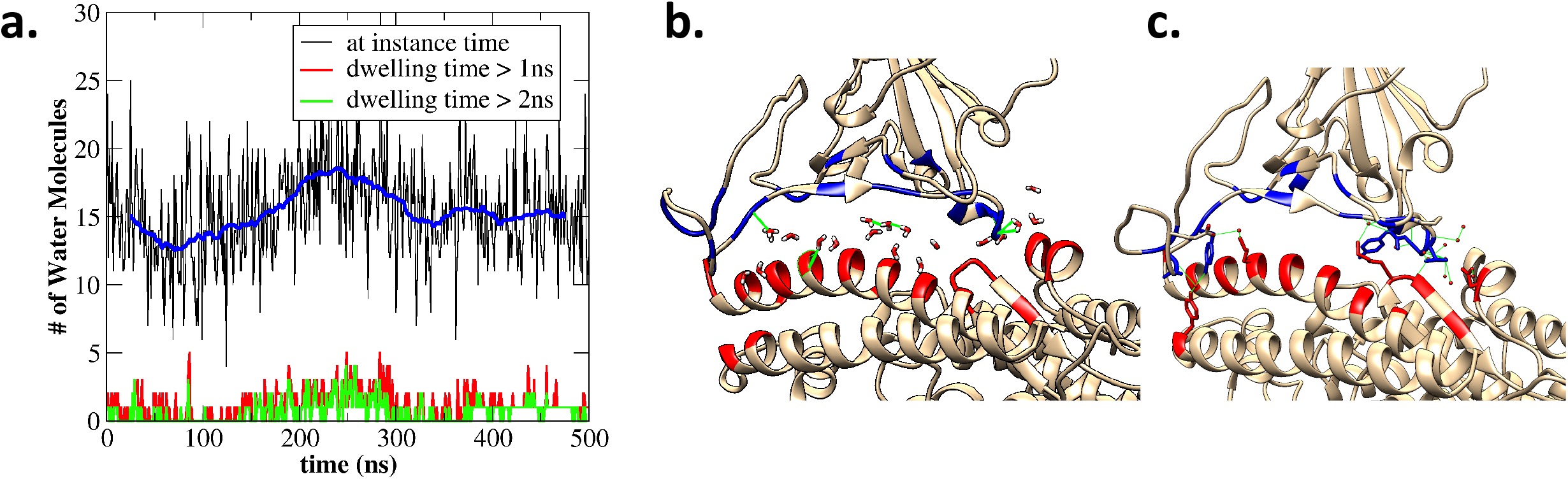
The water molecules at the binding interface. (**a**) The number of water molecules that bridge the CoV2-RBD and ACE2 found in a simulation trajectory. (**b**) The water molecules at the interface for the initial structure after equilibration. (**c**) The water molecules at the interface as observed in the crystal structure.

### 4. Simulation of the human ACE2 mutants uncovers new binding modes

It has been demonstrated that the ACE2 from several mammalian species possess high sequence similarities, yet their binding to the SARS-RBD differs significantly. In particular, the binding of SARS-RBD to the rat ACE2 is much weaker as discovered in experiments^12^ Inspired by these information, two mutants of the CoV2-RBD/ACE2 were constructed: (1) *ACE2-mut-h1* by mutating N-terminal helix-1 to that of the rat ACE2; (2) *ACE2-K353H* by mutating K353 to Histidine (the amino acid in wild type Rat ACE2). Two 500-ns MD simulations were carried out for each mutant system. The simulation showed that the mutations in *ACE2-mut-h1* reduced the interaction between the CoV2-RBD and the helix-1, and the *ACE2-K353H* showed weaker binding between the CoV2-RBD and the β-hairpin centered at the H353. Using the clustering analysis, the representative structures were identified from each simulation trajectory (Figure 7). Although the overall topology is very similar to the wild type complex structure, there are pronounced differences. For the *ACE2-mut-h1* system, the CoV2-RBD tilted further away from the ACE12 helix-1 in one simulation (Figure 7a); and the CoV2-RBD lost its contact with helix-13 (G326 to N330) in another simulation for the *ACE2-K353H* (Figure 7c). In the wild type ACE2, the K353 is a hydrogen donor, and its mutant H353 cannot form the hydrogen bond with the CoV2-RBD as in the wild type CoV2-RBD/ACE2 complex. The number of contacting residue pairs was significantly reduced in the *ACE2-K353H* mutant system. This is in line with the report that K353 is more critical than the other residues, as its hydrophobic neighborhood enables this positively charged residue high selectivity to the RBD^27,28^.

**Figure 7.**
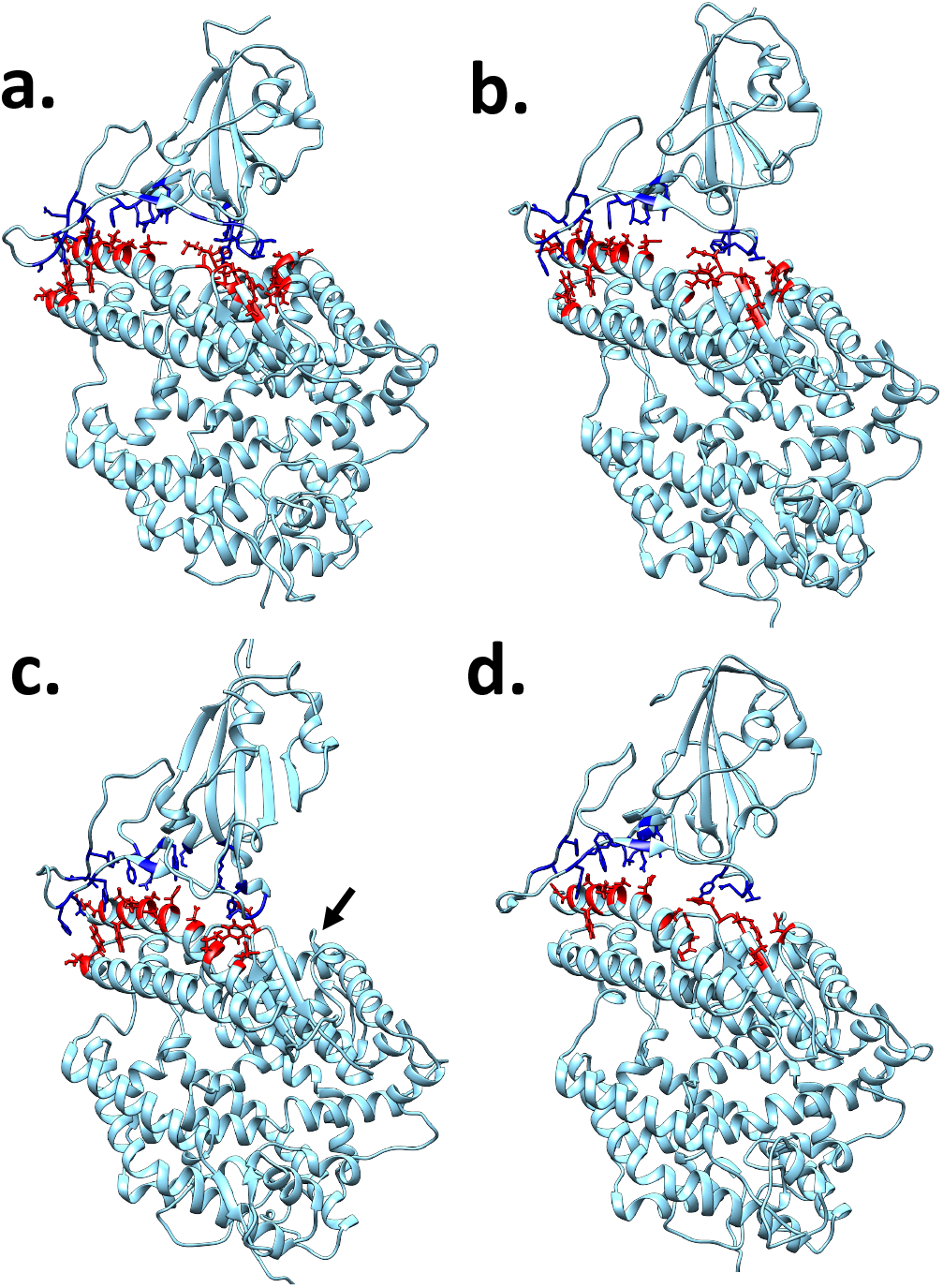
The representative structures for the ACE2-mutants in complex with the CoV2-RBD. (**a, b**) *ACE2-mut-h1*, in which the human ACE2 helix-1 (residues 19-40) was mutated to the Rat sequence. The structure in (a) is the same as the green model shown in Figure 4b, which illustrates a pronounced tilting movement. (**c, d**) *ACE2-K353H,* where the ACE2 K353 was mutated to a histidine. The CoV2-RBD in (c) lost its contact with the Helix-13 of the ACE2 (residues 498-503, indicated by the black arrow). Each structure was obtained by the clustering analysis of 500ns MD simulation trajectories.

The physical interactions between the RBD and the ACE2 were quantified for the simulated structures. We considered the molecular mechanics energy *E_MM_*, which is composed of the van der Waals and the electrostatic interactions. Furthermore, the number of residue contacts *(NC)* between (RBD and ACE2) was extracted from simulated structures. Both the *E_MM_* and *NC* indicate that the RBD interactions with the ACE2 are comparable for CoV2 and SARS spike proteins (Figure 8). From the simulations, the CoV2-RBD has more close contacts with the ACE2 than the SARS-RBD: the numbers of contacts for CoV2-RBD/ACE2 and SARS-RBD/ACE2 are 32.9 and 30.4 respectively. In terms of interaction energies, CoV2-RBD showed slightly weaker interactions with the ACE2 compared to the SARS-RBD (*E_MM_* values are −1017.1 kJ/mol vs. −1133.4 kJ/mol). We would like to point out that the energy E_MM_ is the physical interaction between the RBD and the ACE2, rather than the binding energy, which requires accurately incorporating solvation energy and entropy. Furthermore, the standard deviations of *E_MM_* are 70.2 kJ/mol and 65.5 kJ/mol for the two complexes. Therefore, we infer that the binding affinities are comparable for CoV2-RBD/ACE2 and SARS-RBD/ACE2. The simulations started from the predicted and crystal models yielded very similar results (purple triangles). This is in line with a recent study, in which the authors showed similar binding affinity to human ACE2 for both SARS-CoV-2 and SARS-CoV spike proteins^29^ They found the association rate constants *k_on_* to be the same at 1.4×10^5^ M^-1^s^-1^, while the SARS-CoV spike protein showed a faster dissociation, with the rate constant *k_off_* to be 7.1×10^-4^ s^-1^, about 4.4 times larger than the SARS-CoV-2 spike protein *k_off_* = 1.6×10^-4^ s^-1^. similar *k_on_* values and the equilibrium dissociation constants *K_D_* in nanomolar range were reported in other studies for SARS-CoV-2 spike protein (or RBD) binding to human ACE2^10,30^.

**Figure 8.**
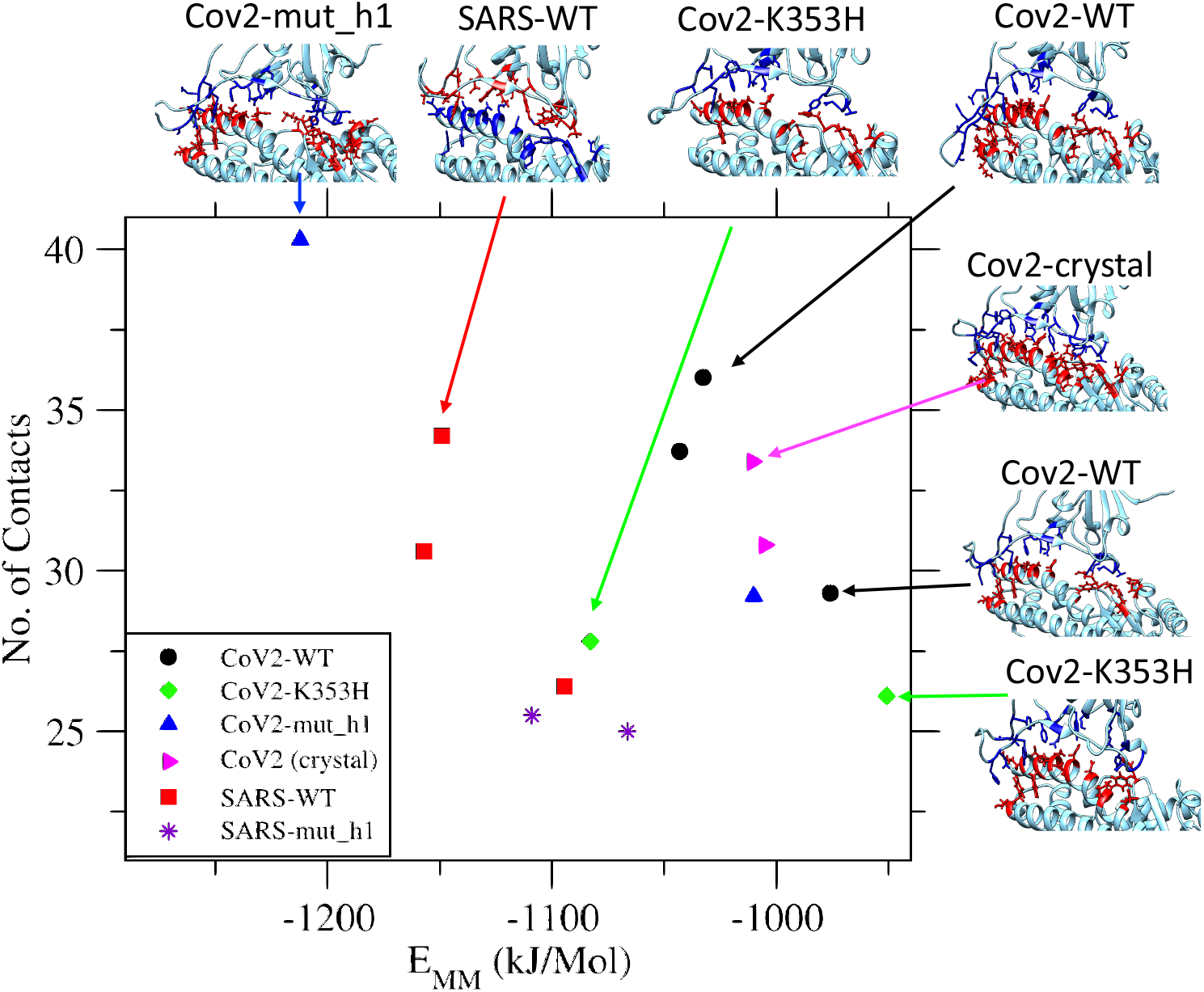
The interaction energies and the residue contacts between the RBD and the ACE2. The molecular mechanics interactions and the number of contacting residue pairs were obtained from the structures sampled by MD simulations. In the legend, the CoV2 and SARS indicate the sources of the RBD; the WT, K353H, mut_h1 are the abbreviations for different constructs of the human ACE2, corresponding to the wild type, the mut_h1, and the K353H.

More interestingly, the mutation impacts were reflected in the *E_MM_* and *NC* analysis: the *ACE2-mut_h1* is likely to reduce the binding to the ACE2 due to the tilting movement of CoV2-RBD, making it further from the ACE2 helix-1 (the blue triangle symbol at lower right, see Figure 7a for the representative structure). In the other simulation trajectory for the CoV2-RBD/ACE2-mut_h1 complex (blue triangle at the left upper corner), the largest *NC* was observed among all simulations. For simulations of the complex with *ACE2-K353H* mutants (green diamonds), the number of contacts were both reduced compared to the wild type system. In one simulation, the contacts between the CoV2-RBD and the Helix-13 of the ACE2 were completely lost (see Figure 7c), consistent with the less favorable interactions reflected on an increase of *E_MM_*. For the SARS-RBD interaction with the *ACE2-mut_h1*, both simulations revealed fewer contacts compared to the wild type SARS-RBD/ACE2 complex (purple stars in Figure 8). Although direct observation of dissociation is beyond the reach of all-atom MD simulations, the reduced contacts for mutant ACE2 indicates the weaker binding affinity compared to the wild type human ACE2.

## Discussions and Conclusion

The homology modeling of the CoV2-RBD/ACE2 complex yielded highly consistent models compared to the crystal structures. All-atom molecular dynamics simulations were carried out to study the dynamic interactions of CoV2-RBD with human ACE2, the results were compared to the SARS-RBD/ACE2 system. The human ACE2 mutants were also constructed to mimic the rat ACE2 to investigate the roles of critical residues, and possible binding modes in other mammals. It is observed that MD simulations improved the structure at the binding interface and strengthened the interactions between the subunits. The structure of the complex interface is highly stable for all simulations of CoV2-RBD/ACE2 complex in the wild type. The loop region between β6 and β7 can potentially form more contacts with the ACE2 as observed in one simulation trajectory. The simulations results also reveal that the interactions between CoV2-RBD and the ACE2 are mediated by water molecules at the interfaces, stressing the necessity of accounting for the explicit water molecules when quantifying the binding affinity.

The interactions between the RBD and the ACE2 were quantified by physical interaction energies (molecular mechanics energy) and the number of contacting residues. The detailed comparison results suggest that the CoV2-RBD and the SARS-RBD bind to human ACE2 with comparable affinity. The comparison between the SARS-RBD/ACE2 and the CoV2-RBD/ACE2 complexes, with the former forms fewer contacts than the latter (Figure 8), yet exhibiting stronger interactions. The decomposition of the *E_MM_* to the van der Waals and the electrostatic interactions revealed that the major difference is attributed to the electrostatic interactions. Furthermore, we compared the major contacting residues and found that the SARS-RBD has two charged residues (R426 and D463) and the CoV2-RBD has only one charged residue (K417) at the complex interface. The polar and hydrophobic residues are comparable in the two RBDs. This is consistent with the statistics of hydrogen bonds at the complex interfaces.

This study was started with a structure predicted using homology modeling method, which later found to be highly consistent with the crystal structure, demonstrating the potentiality of structure prediction and dynamics simulation in revealing molecular details before the availability of high resolution experimental information. Furthermore, the interactions between CoV2-RBD and the ACE2 mutants mimicking rat ACE2 protein were investigated. The results provide valuable information at the atomic level for the reduced binding affinity. The recent report on the SARS-CoV-2 infection to a dog raise the concerns about the COVID-19 transmissions to pet animals. The approach discussed in this study can be applied to analyze the CoV2-RBD interaction with a broader ACE2 variants, including the genetic polymorphism of the ACE2 in human and the ACE2 of other mammalian animals. Preliminary analysis of several ACE2 proteins from mammalian animals using homology modeling and active site prediction methods shows that the binding sites exhibit different features compared to the human ACE2. More thorough and deeper analysis can be carried out using the presented approach.

The high quality homology model allowed us to start the simulation analysis before the high resolution experimental data became available. The protein structure prediction community are making efforts to provide high quality models using advanced modeling and prediction methods^32–35^. Based on predicted and experimentally determined structures, molecular dynamics simulations will add more insights to the molecular mechanism and allow better probing properties due to mutation or modification to proteins.

## Acknowledgement

The work is supported by Beijing Computational Science Research Center (CSRC) via a director discretionary grant. The authors acknowledge the Beijing Super Cloud Computing Center (BSCC) for providing HPC resources that have contributed to the research.

