## supplementary information file for "Computational simulations reveal the binding dynamics between human ACE2 and the receptor binding domain of SARS-CoV-2 spike protein"

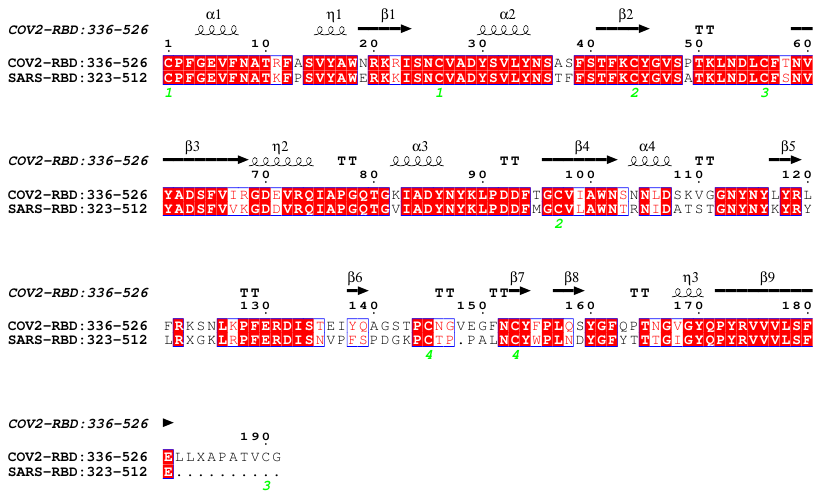

**Figure S1. The sequence alignment of receptor binding domain (RBD) for SARS-COV-2 and SARS-COV.**

**a.**

With respect to starting model

With respect to representative model

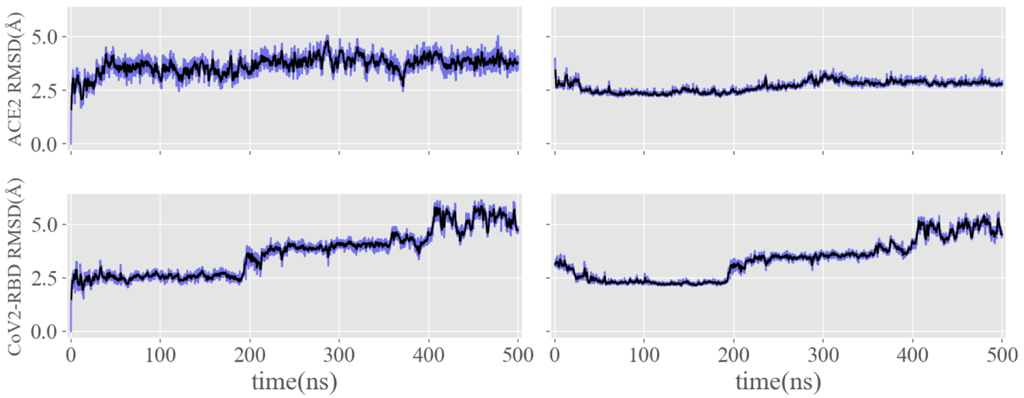

**b.**

With respect to representative model

With respect to starting model

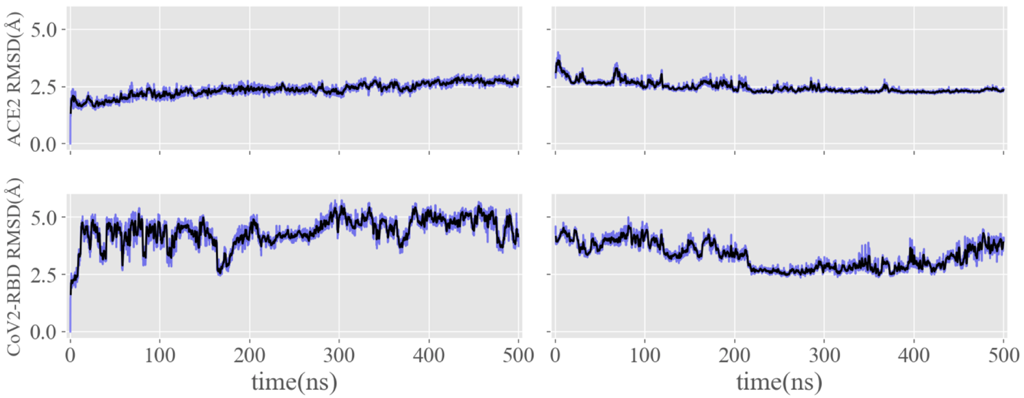

**Figure S2. The structure difference compared to the starting and representative models.** (**a**) simulation trajectory #2; (**b**) simulation trajectory #3.

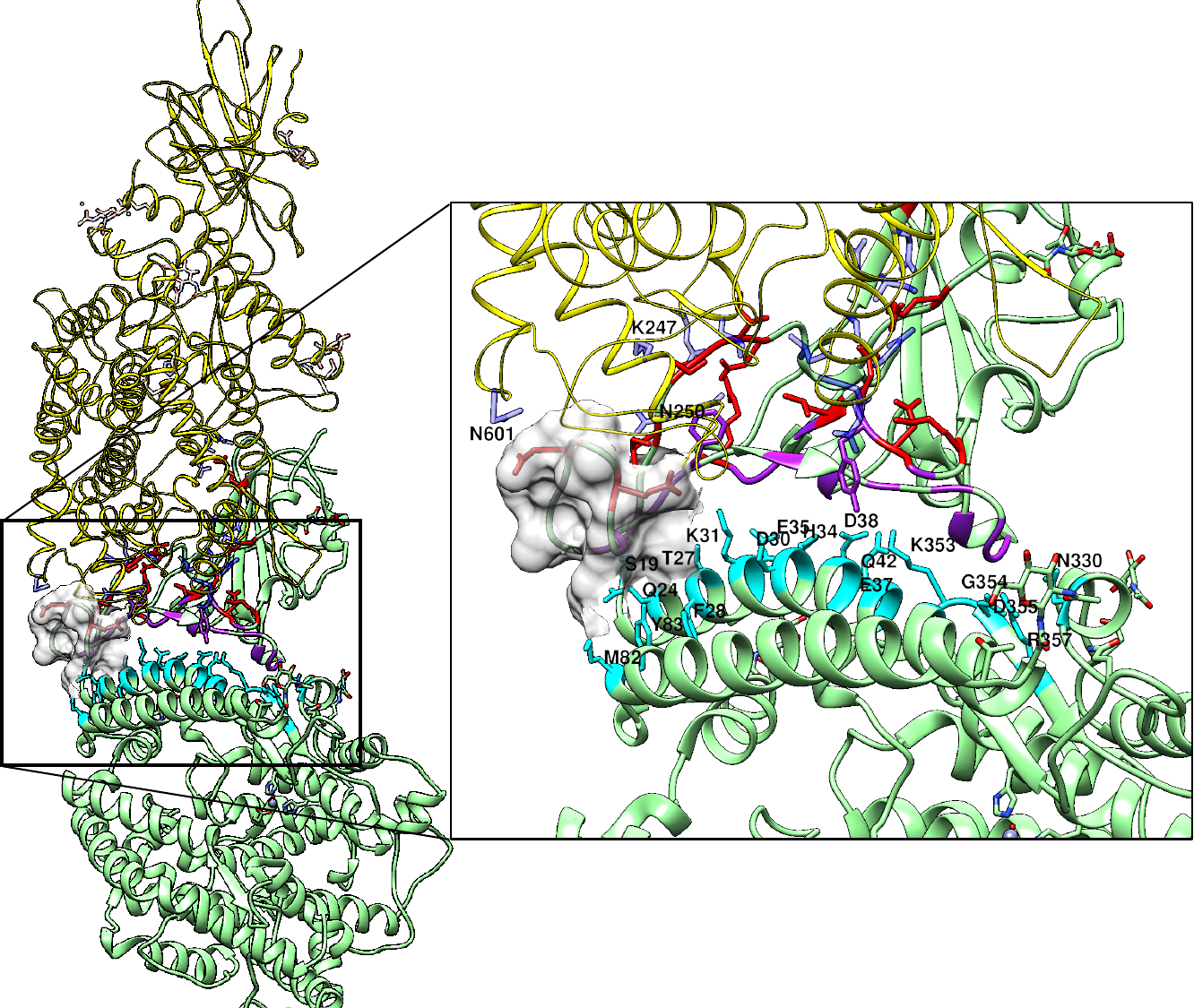

**Figure S3.** **The contacts between CoV2-RBD and the ACE2 in crystal.** Two copies of the CoV2-RBD/ACE2 complex (asymmetric unit) are shown in green and yellow color. The loop region between β6 and β7 of the CoV2-RBD is highlighted with the surface representation. This region interacts with two ACE2 molecules: one at the binding site (green), and the other one (yellow) is due to the crystal packing. The cyan color highlights the contacting residues of ACE2.

**a.**

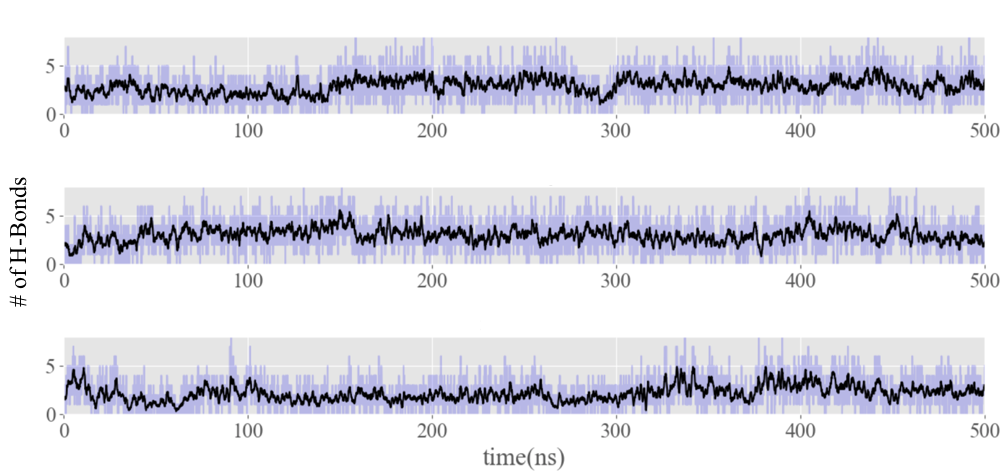

**b.**

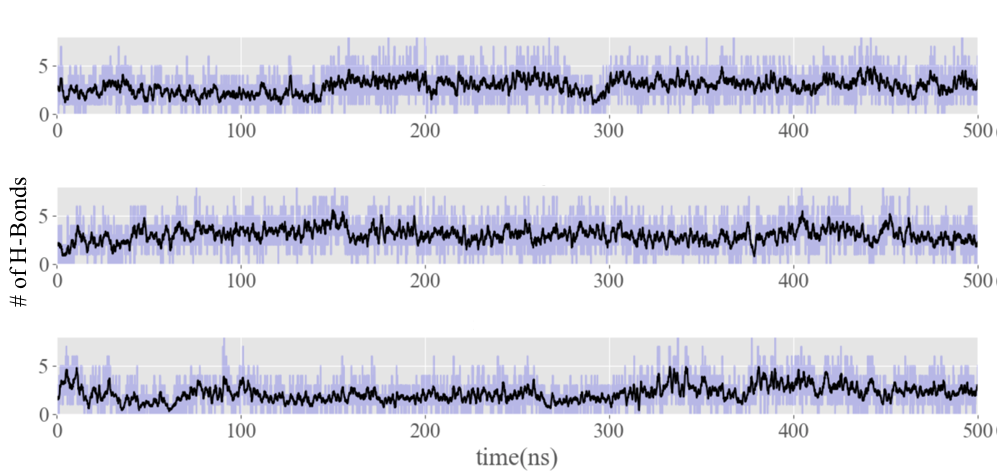

**Figure S4. The number of hydrogen bonds at the complex interface using the default donor-receptor distance cutoff=3.0 Å.** (**a**) hydrogen bonds for CoV2-RBD/ACE2 complex; (**b**) hydrogen bonds for SARS-RBD/ACE2 complex.

| Table S1. Contact residues at the SARS-COV- RBD/ACE2 interface | | | | | | |
| --- | --- | --- | --- | --- | --- | --- |
| Traj 1 | | | **Traj2** | | **Traj 3** | |
| *ACE2* | | *CoV* | *ACE2* | *CoV* | *ACE2* | *CoV* |
| S19  Q24  T27  K31  H34  E37  D38  Y41  Q42  L45L  L79L  M82  Y83  Q325  E329  N330  K353  G354  D355  R357 | R426  Y436  Y440  Y442 L443  D463 L472  N473  Y475  N479  G482  Y484 T486 T487  G488 I489 Y491 | | Q24  T27  D30  K31  H34  E37  D38  Y41  Q42  L45L  L79L  M82  Y83  Q325  E329  N330  K353  G354  D355  R357 | R426  Y436  Y440  Y442 L443  P462  D463  G464 L472  N473  Y475  N479  G482  Y484 T486 T487  G488 I489 Y491 | Q24  T27  K31  H34  E37  D38  Y41  Q42  L45L  L79L  M82  Y83  Q325  E329  N330  K353  G354  D355  R357 | R426  Y440  Y442 L443  P462  D463  G464  P470 L472  N473  Y475  L478  N479  G482  Y484 T486 T487  G488 I489 Y491 |
